# The CRISPRaTOR: a biomolecular circuit for Automatic Gene Regulation in Mammalian Cells with CRISPR technology

**DOI:** 10.1101/2024.03.30.587417

**Authors:** Alessio Mallozzi, Virginia Fusco, Francesco Ragazzini, Diego di Bernardo

## Abstract

We introduce the CRISPRaTOR, a biomolecular circuit for precise control of gene expression in mammalian cells. The CRISPRaTOR leverages the stochiometric interaction between the artificial transcription factor VPR-dCas9, and the anti-CRISPR protein AcrIIA4, enhanced with synthetic coiled-coil domains to boost their interaction, to maintain the expression of a reporter protein constant across diverse experimental conditions, including fluctuations in protein degradation rates and plasmid concentrations, by automatically adjusting its mRNA level. This capability, known as Robust Perfect Adaptation (RPA), is crucial for the stable functioning of biological systems and has wide-ranging implications for biotechnological applications. The CRISPRaTOR belongs to a class of biomolecular circuits named antithetic integral controllers, and it can be easily adapted to regulate any endogenous transcription factor thanks to the versatility of CRISPR-Cas system. Finally, we show that RPA holds also in cells genomically integrated with the CRISPRaTOR, thus paving the way for practical applications in biotechnology that require stable cell lines.

## Introduction

Robustness is a fundamental property that enables biological systems to maintain stability and functionality despite fluctuations in biochemical reaction rates caused by environmental perturbations. Robust Perfect Adaptation (RPA) is a specific form of robustness in which a biological system returns to a baseline level of function despite external fluctuations, providing a mechanism for homeostasis^1^. RPA mechanisms are often found in cellular signalling pathways, where they help maintain a consistent response to signals despite changes in signal intensity. Well-studied examples of RPA in biomolecular processes are calcium homeostasis within mammalian cells^2^ and the chemotaxis in E. Coli^3,4^, allowing organisms to adapt to changing concentrations of nutrients, toxins, or hormones.

Control Engineering is a well-established discipline to build “controllers” to regulate the behavior of a physical system by keeping its output constant across a range of operating conditions, by dynamically adjusting the input. In the context of biological processes, the ***input*** can be any molecular species (e.g. a small molecule, a metabolite, etc.) whose changes have a measurable effect on the ***output*** of the biological process (e.g. a protein of interest).

A key theoretical result of Control Engineering is that negative feedback can endow systems with robustness to perturbations and uncertainties^5^. Specifically, a negative-feedback controller relies on a sense and react paradigm, where the output is actively measured and compared against a reference value. Depending on the difference between the two (control error), the controller will dynamically adjust the input to minimize the control error. It can be demonstrated that, under specific conditions, if the magnitude of the input is proportional to the sum of the error over time (integral control), then the system will exhibit RPA^6,7^.

Thanks to recent advances in molecular biology and biomolecular control theory, building a biomolecular integral controller to robustly regulate gene expression at a constant level has now become feasible. A biomolecular implementation of an integral controller is shown in **Figure 1**, and it has been named the Antithetic Integral Controller (AIC)^8^. It consists of an activator *(X)*, which drives the expression of the “output” species *(Z)*, and an inhibitor *(Y)*, which stoichiometrically binds *X* in a one-to-one fashion and inactivates it. In this configuration, if a decrease in *Z* occurs, because of an external perturbation, it will cause a decrease in Y and thus free up more *X* to increase the level of *Z* back to its initial level. Similarly, an increase in *Z* will indirectly decrease *X* via *Y* and thus reestablish the equilibrium.

**Figure 1.**
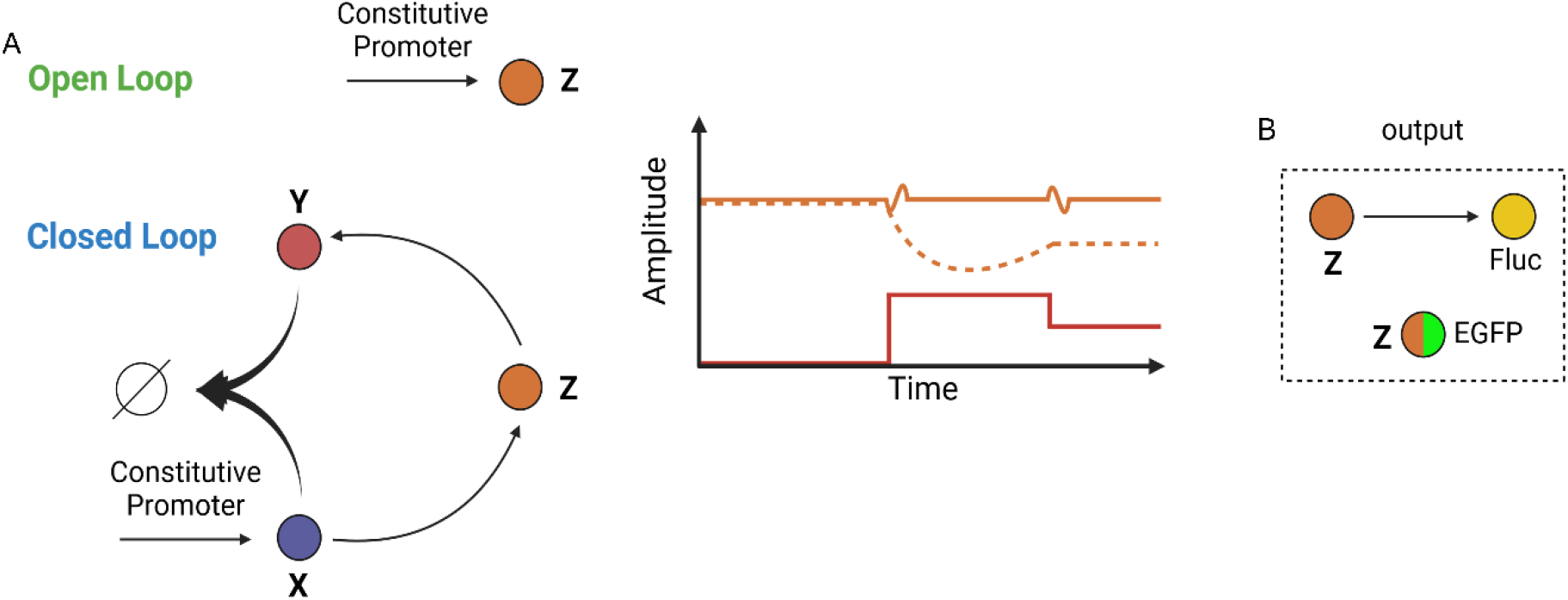
The Antithetic Integral Controller and Robust Perfect Adaptation. **A)** The Antithetic Integral Controller is a negative feedback loop (Closed Loop) where a constitutively expressed activator species X drives the expression of a species of interest Z (output). Z drives the expression of an inhibitor species Y, which binds and inhibits X. When the concentration of Z changes, so does Y thus causing X to change in an opposite manner to Z (e.g. if the concentration of Z decreases, active X will increase, and vice versa). This mechanism enables the Antithetic Integral Controller to dynamically adjust the concentration of Z (solid orange line) in the face of a perturbation (red line) and thus maintain Z constant over time. In the Open Loop configuration, Z is directly expressed from a constitutive promoter, and if its concentration decreases because of an external perturbation (red line), its concentration would not be constant over time (dashed orange line). **B)**The species Z in our implementation is itself a transcriptional activator and its concentration can be experimentally tracked over time either indirectly by placing the luminescence firefly Luciferase (Fluc) under a promoter driven by Z, or directly by fusing the EGFP fluorescent reporter to Z itself.

The AIC has been experimentally implemented in bacteria using sigma/anti-sigma factors in bacteria^9^, and more recently in mammalian cells by means of a pair of sense and antisense mRNA^10^, or by protein splicing with inteins^11^. These implementations albeit successful, have some limitations in mammalian systems: the use of sense-antisense RNA pairs could potentially trigger toxicity because of the cell’s innate immune response, as double-stranded RNAs are generated during viral replication^12^; the protein splicing approach, despite being potentially compatible to any protein of interest, requires extensive engineering to ensure correct inteins splicing and protein folding. Furthermore, these AIC implementations have been tested by transient transfection, where plasmid molar ratio can be strictly controlled. However, engineering cells for practical application would require stable integration of gene circuits in the cells’ own genome, and proof that performances would still be the same are lacking.

Here, we leveraged the proteins of the CRISPR/Cas family to implement an AIC in mammalian cells. Specifically, we made use of the artificial transcription factor VPR-dCas9^13^ and the recently discovered anti-CRISPR protein AcrIIA4^14,15^, augmented with synthetic coiled-coil domains to enhance their binding affinity^16,17^, to implement an AIC in mammalian cells. We demonstrate its ability to confer RPA in both transient transfection and stable genomic integration. The use of VPR-dCas9 makes our implementation very versatile, as it can be used as a “plug-and-play” circuit to control any endogenous transcription factor of interest by simply designing an appropriate guide RNA (gRNA). We named our implementation of the AIC, the CRISPRaTOR.

## Results and Discussion

### Experimental implementation of an Antithetic Integral Controller (AIC) by means of the CRISPR-antiCRISPR system

The experimental implementation of the AIC is shown in **Figure 2**. The nuclease-deficient Cas9 fused to the transactivation domains VP64, p65, and Rta (VPR) (VPR-dCas9)^13^ acts as species **X** (**Figure 1**) and it is constitutively expressed from the pCMV promoter. This synthetic transcriptional activator can drive transcription from any synthetic or endogenous promoter of interest by simply choosing the cognate guide-RNA (gRNA). As species **Z** of the AIC (**Figure 1**), we chose the Reverse Tet TransActivator (rtTA) driven by the 7B_pMin promoter^18^, which in the presence of the constitutively expressed gRNA_B, is activated by the VPR-dCas9-N8. In the presence of Doxycycline, rtTA binds the pTRE3G promoter upstream of the anti-CRISPR protein AcrIIA4, which acts as species **Y** since it can bind and inactivate the cognate Cas9 ^14,15^. We have previously demonstrated^19^ that the inhibitory activity of AcrIIA4 towards the VPR-dCas9 can be increased by more the 3-fold by boosting their binding affinity thanks to the fusion of two orthogonal synthetic Coiled Coils (CCs) domains (N7 and N8^16,17^) thus giving rise to two new moieties, namely AcrIIA4-N7 and VPR-dCas9-N8, as shown in **Figure 2A**. Our AIC implementation, which we named the CRISPRaTOR, thus exploits the stoichiometric inhibitory action of AcrIIA4-N7 to negatively regulate the transactivator VPR-dCas9-N8 and to give rise to a negative feedback regulation of the rtTA protein and to Robust Perfect Adaptation (RPA). For example, if the rtTA protein level decreases, then there will be less transcription from the pTREG promoter, and hence less AcrIIA4-N7 protein. This in turn will lead to an increase in “free” VPR-dCas9-N8 (i.e. not bound by AcrIIA4-N7) and consequently an increase in rtTA transcription, thus eventually re-equilibrating the level of rtTA. To monitor the level of rtTA, we used two alternative strategies as summarised in **Figure 1**, and detailed in **Figure 2A** and **Figure 3A**. Specifically, we either cloned the firefly luciferase (Fluc) downstream of a pTRE3G promoter, so that by measuring luminescence we have an indirect readout of rtTA level (**Figure 2A**), or alternatively we fused a fluorescence tag to the rtTA and we monitored green fluorescence levels (**Figure 3A**).

**Figure 2.**
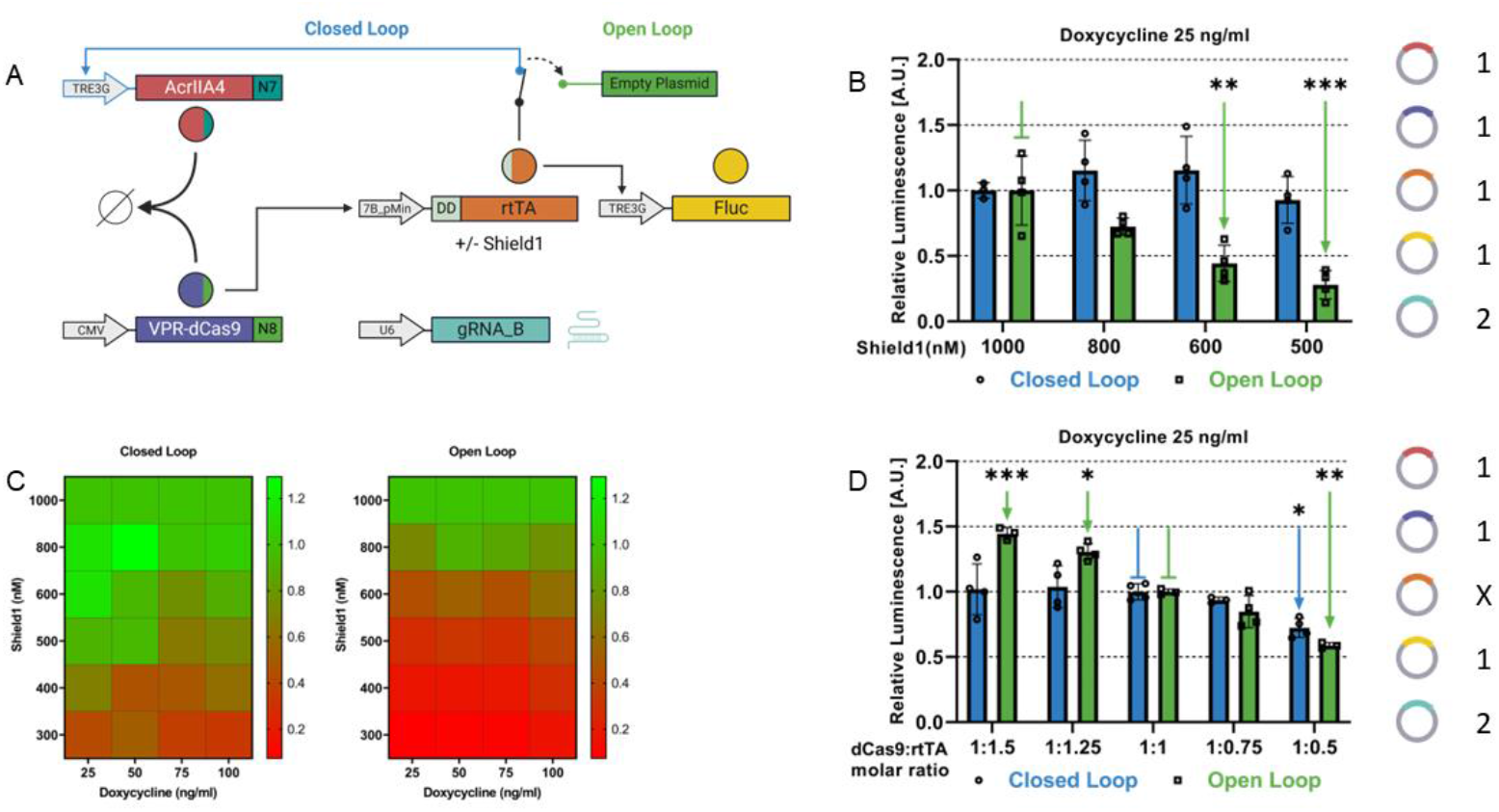
The CRISPRaTOR, an antithetic integral controller based on the CRISPR-antiCRIPSR system: **A)** Schematic representation of the CRISPRaTOR. The DD-rtTA drives the expression of the Fluc from the pTRE3G promoter, and, only in the Closed Loop configuration, also of the antiCRISPR AcrIIA4-N7. In the Open Loop configuration, the plasmid encoding for the anti-CRISPR protein is substituted by an empty plasmid. **B)** Experimental validation of robust perfect adaptation to changes in protein degradation. Relative Fluc luminescence is computed as the Fluc luminescence first normalized to the Renilla luminescence at the indicated concentration of the Shield1 molecule, and then divided by its value at 1000 nM of Shield1. Doxycycline is kept constant at 25ng/ml. The green pointed arrows indicate a significant difference in relative luminescence versus the value indicated by the green blunted arrow. Transfected plasmids with molar ratios are schematically represented as colored circles with numbers indicating relative molar ratios. **C)** Experimental exploration of the parameter space in which RPA is achieved. The heatmaps report the relative Fluc luminescence at the indicated concentration of Shield1 and Doxycycline for the CRISPRaTOR in Closed Loop and Open Loop configurations. **D)** Experimental validation of robust perfect adaptation to changes in plasmid ratios. Relative Fluc luminescence values are computed as the normalized Fluc luminescence at the indicated molar ratios divided by its value at the 1:1 molar ratio. Transfected plasmids with molar ratios are schematically represented as colored circles with numbers indicating relative molar ratios, while the X indicates the plasmid whose ratio is being changed. The concentration of doxycycline is kept constant at 25ng/ml. Shield1 is kept constant at 1000nM to stabilize DD-rtTA. The pointed arrows indicate a significant difference in relative luminescence versus the value indicated by the blunted arrow of the same color (blue or green). ***VPR-dCas9-N8***: *nuclease-deficient Cas9 fused to the transactivation domain VPR and to synthetic coiled-coil N8;* ***AcrIIA4-N7***: *Anti-CRISPR protein fused to the synthetic coiled-coil N7;* ***DD-rtTA***: *reverse tetracycline TransActivator 3G fused to the FKBP derived Destabilization Domain (DD) whose degradation is modulated by the small molecule Shield1*.; ***TRE3G:*** *Tetracycline Responsive Element promoter 3G*. ***gRNA_B***: *gude RNA with sequence B;* ***FLuc***: *firefly Luciferase. n=4 biological replicates. A minimum of n=3 when one of the measurements was identified as an outlier (Grubbs’ test, alpha=0*.*2). Statistics analysis has been conducted through a two-way ANOVA test. * P ≤ 0*.*05 ** P ≤ 0*.*01 *** P ≤ 0*.*001 **** P ≤ 0*.*0001*.

**Figure 3.**
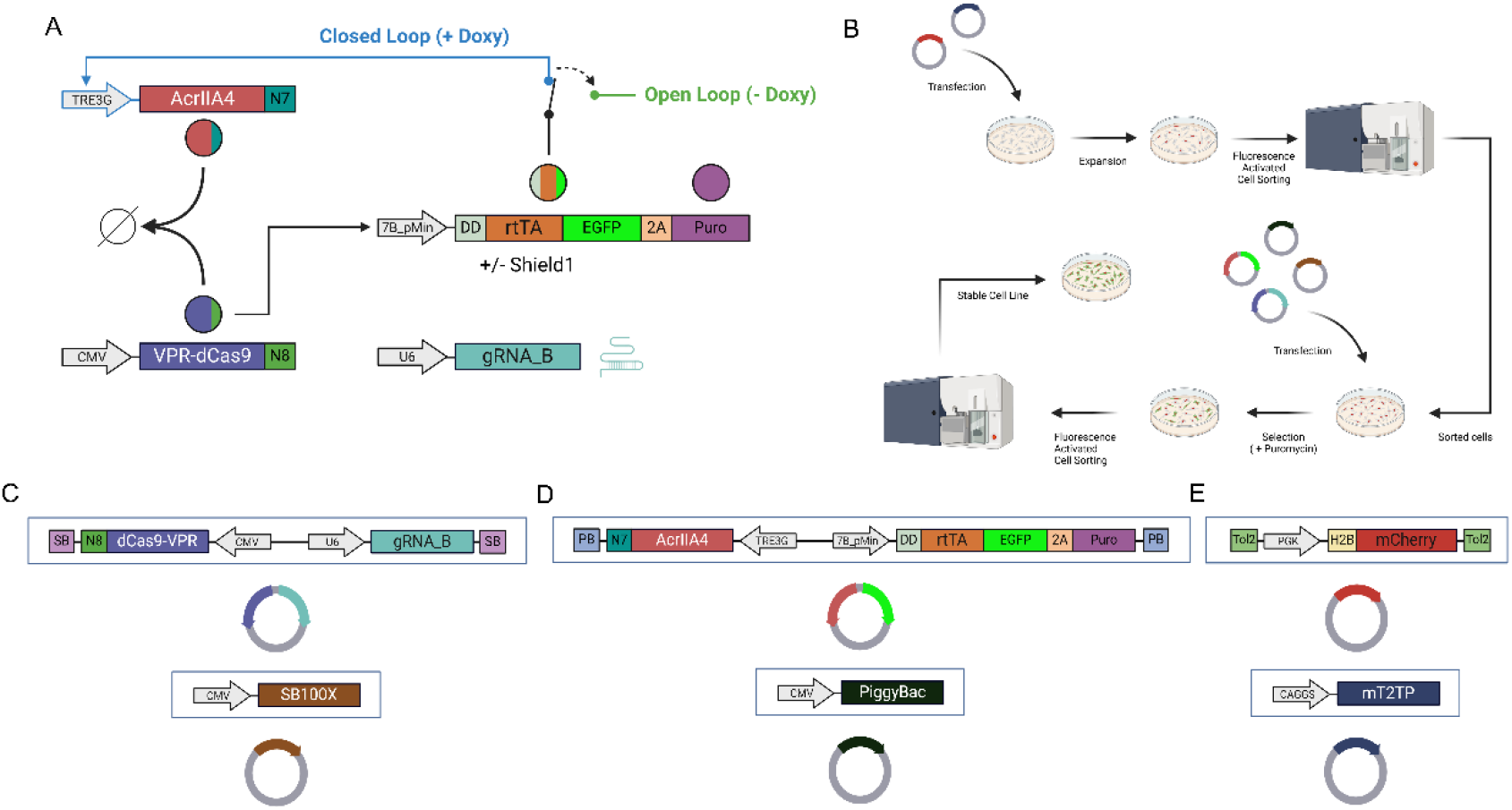
Development of a cell line with stable integration of the CRISPARATOR: **A)** Schematic representation of the CRISPARATOR genomically integrated in Hek293T cells. The destabilization domain (DD) is fused at the N-terminal of the rtTA protein, whereas at its C-terminal, the fluorescent protein EGFP and the Puromycin resistance protein separated by the self-cleaving 2A peptide are found. The circuit exhibits two distinct operational states: the Closed Loop and the Open Loop. The Closed Loop configuration is achieved upon the addition of Doxycycline, when the AcrIIA4 protein is expressed. Conversely, in the absence of Doxycycline, the system transitions to the Open Loop state, where the AcrIIA4 protein cannot be expressed. **B)** Schematic representation of the strategy to genomically integrate the CRISPRaTOR by means of transposases. The H2B_mCherry is integrated with the Tol2 transposase through multiple rounds of cell sorting post transfection. Once a stable mCherry-expressing cell line is obtained, constructs codifying for the CRISPRaTOR are transfected along with their transposases. Puromycin selection occurs only when both plasmids encoding for the CRISPARATOR are present. Sorting for mCherry and EGFP are subsequently performed as quality control. **C)** Plasmid codifying for the VPR-dCas9_N8, under the control of the strong, constitutive CMV promoter and the guide-RNA gRNA_B under the control of the U6 promoter. The SleepingBeauty transposase is on a second plasmid under the control of the CMV promoter and recognizes the SB sequences **D)** Plasmid codifying for the AcrIIA4-N7, under the control of the Doxycycline-inducible TRE3G promoter, and for the DD-rtTA-EGFP-2A-Puromycin, under the control of 7B_pMin promoter. On the second plasmid, the PiggyBac transposase, under the control of the CMV promoter, recognizes the PB sequence. **E)** Plasmid codifying for H2B_mCherry, under the control of the weak, constitutive promoter PGK, and a second plasmid codifying for the transposase mT2TP, under the control of the CAGGS promoter. The mT2TP transposase recognizes the Tol2 sequences.

**Figure 4.**
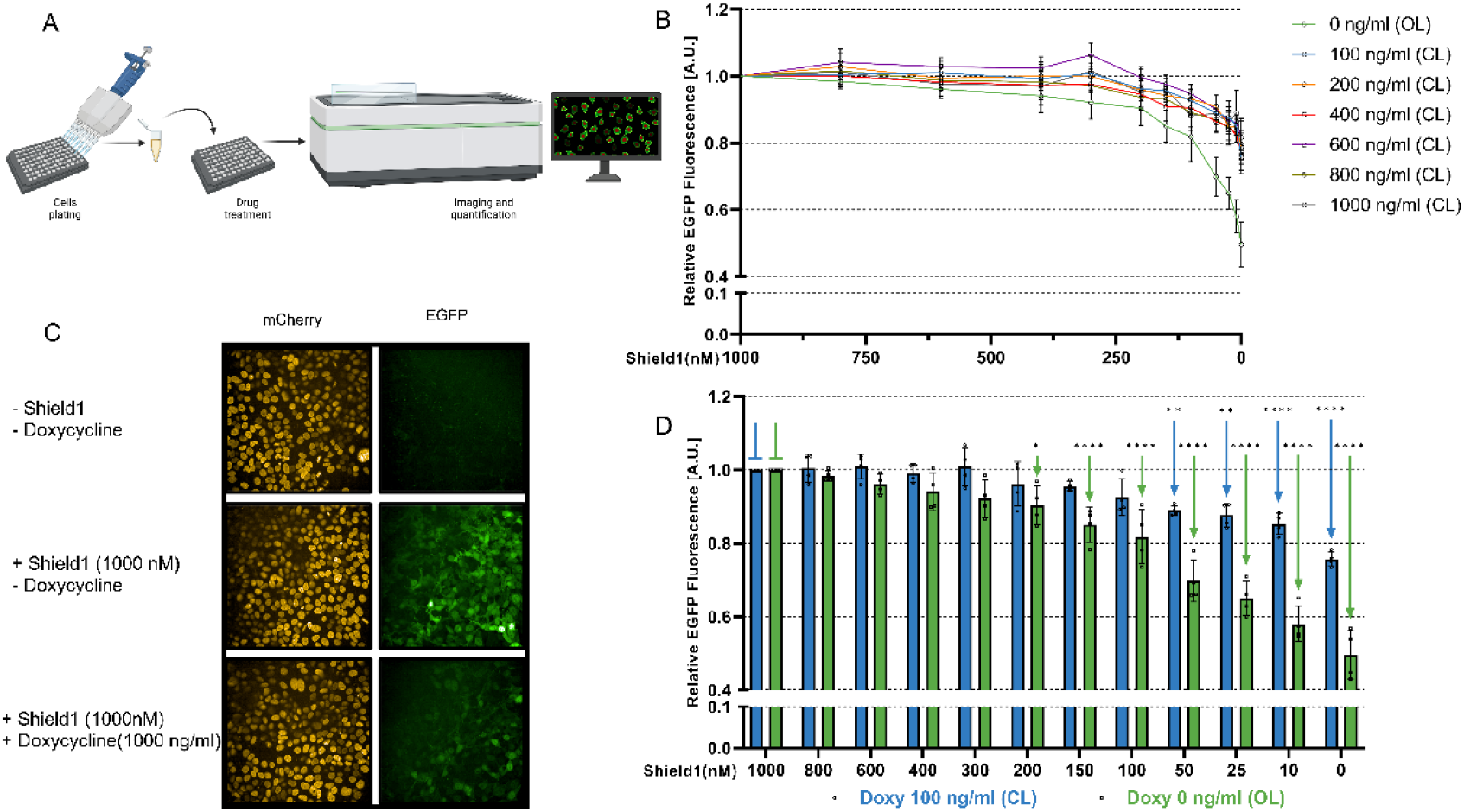
Robust Perfect Adaptation in stable cell lines: **A)** Schematic representation of the workflow of the High Content Screening platform used to detect and quantify EGFP fluorescence to track rtTA levels in CRISPRaTOR-containing Hek293T cells. Cells were plated in a 96-well plate and, 18 hours after seeding, treated with different concentrations of Shield1 and Doxycycline. Forty-eight hours after treatment, cells were imaged using the Opera Phenix. **B)** Quantification of the green fluorescent signal from confocal images in Hek293T cells integrated with the CRISPRaTOR for decreasing concentrations of Shield1 and for the indicated fixed concentration of Doxycycline. In the absence of Doxycycline, the CRISPRaTOR is in the Open Loop configuration. Fluorescence values are relative to the value measured at 1000 nM Shield1. **C)** Representative fluorescence images at confocal microscope of Hek293T cells integrated with the CRISPRaTOR following the indicated treatments. **D)** Relative fluorescence measured as in (C) but represented as a bar plot for only two conditions: Doxycycle 100 ng/ml (blue) and without Doxycycline (green). The pointed arrows indicate a significant difference in relative luminescence versus the value indicated by the blunted arrow of the same color (blue or green). *For imaging quantification experiments, n=4 biological replicates coming from 4 96 well plates. Statistics analysis has been conducted through a two-way ANOVA test. * P ≤ 0*.*05 ** P ≤ 0*.*01 *** P ≤ 0*.*001 **** P ≤ 0*.*0001*.

### The CRISPRaTOR confers Robust Perfect Adaptation in transient transfection

To test RPA, we fused the bacterial derived Destabilization Domain (DD)^20^, a 12-kDa (107-amino-acid) tag based on a mutated FKBP, to the transcription factor rtTA to obtain DD-rtTA, as shown in **Figure 2A**. By changing the concentration of the small molecule Shield1, it is possible to change the stability of the DD-rtTA fusion protein and thus its amount. We first assessed the degradation of DD-rtTA protein and its rescue by Shield1 while also confirming that the DD-rtTA fusion protein still maintains its function as a transcriptional activator. To this end, as shown in **Supplementary Fig. 1A**, we cloned the mCherry reporter gene downstream of the pTRE3G promoter while constitutively expressing the DD-rtTA from the pEF1α promoter. We then measured the fluorescence level in individual cells by flow cytometry in the presence or absence of Shield1 and Doxycycline, as shown in **Supplementary Fig. 1B, 1C**.

As Shield1 stabilizes the DD-rtTA protein, while Doxycycline allows it to bind to the pTRE3G promoter, then full expression of mCherry is achievable only in the presence of both drugs, as confirmed by our experimental results.

Having confirmed that DD-rtTA acts as a bona-fidae transcriptional activator whose level can be modulated by Shield1, we tested the ability of the CRISPRaTOR in **Figure 2A** in keeping the expression of the Fluc from the pTRE3G promoter stable in the face of changes in Shield1 concentrations, starting at a saturating dose of 1000 nM, which stabilizes the DD-tagged protein. As a negative control, we implemented an *Open Loop* circuit where AcrIIA4-N7 is absent and substituted by an empty plasmid. Since the CRISPRaTOR is encoded on five different plasmids as shown in **Supplementary Fig. 1D**, we performed the experiment in transient transfection in Hek293T cells by transfecting equimolar ratios of all the plasmids except for the one encoding the guide-RNA, whose molar concentration was doubled. As reported in **Fig. 2B and Supplementary Fig. 2A**, for a fixed concentration of Doxycycline of 25 ng/ml, the Fluc luminescence in the *Open Loop* circuit decreases proportionally with the Shield1 concentration. This is expected, as the DD-rtTA protein stability decreases with decreasing concentration of Shield 1 while its mRNA level is unchanged, thus causing an overall decrease in DD-rtTA protein level at equilibrium, and hence of the Fluc expression downstream of the pTRE3G promoter. On the contrary, in the case of the CRISPRaTOR (**Fig. 2B** - blue), the luminescence level remains constant, thus demonstrating Robust Perfect Adaptation; this can be explained by the fact that as DD-rtTA protein stability decreases, this transiently decreases the DD-rtTA protein level, which in turn decreases expression of antiCRISPR, and frees up more VPR-dCas9-N8 transactivator and consequently increased transcription of the *DD-rtTA* mRNA, thus re-establishing the correct level of DD-rtTA protein and hence of Fluc expression. To further explore the experimental conditions in which Robust Perfect Adaptation is maintained, we repeated the same experiment, but fixing Doxycycline concentrations at either 50 ng/ml, 75 ng/ml, or 100 ng/ml, and changing Shield1 concentrations from 300nM to 1000nM. The results are summarised in **Figure 2C** and **Supplementary Fig 2B-E**. The overall results demonstrate that only the closed loop CRISPRaTOR is able to maintain Fluc expression stable across a wide range of experimental conditions.

To further investigate the ability of the CRISPRaTOR in maintaining the FLuc expression stable in the context of transient transfection, we changed the amount of the plasmid encoding DD-rtTA relative two the other two plasmids encoding VPR-dCas9-N8 and AcrIIA4-N7, to assess the ability of the CRISPRaTOR to counteract plasmid copy number variations. As shown in **Fig. 2E and Supplementary Fig. 2F**, increasing or decreasing the relative amount of the DD-rtTA up to 50% had a strong effect on the open loop circuit, on the contrary the CRISPRaTOR was able to maintain the relative luminescence constant for most of the plasmid concentrations. Taken together these results demonstrate that the CRISPRaTOR implements an effective AIC motif conferring Robust Perfect Adaptation.

### Cells with stable integration of the CRISPRaTOR

Upon successful characterization of the biomolecular circuit through transient transfection, we aimed at testing the CRISPRaTOR in a more physiologically relevant condition that could be encountered in biotechnological and therapeutic applications. Hence, we generated a cell line with stable genomic integration of the CRISPRaTOR to verify whether Robust Perfect Adaptation would hold also in this setting.

To facilitate integration, we reduced the size of the CRISPRaTOR while maintaining the ability to assess its robustness, by modifying it as reported in **Figure 3A**. Specifically, we chose to directly monitor the DD-rtTA protein expression by fusing the EGFP fluorescent protein to its C-terminus. Additionally, we employed a P2A sequence to co-express the Puromycin resistance protein from the same construct for subsequent selection of stably integrated cells. We then encoded the CRISPRaTOR on only two plasmids as reported in **Figure 3C, D**. To prevent promoter crosstalk and independent expression of each cistron^21^, we cloned the two expression cassettes in each plasmid in reverse orientation. To genomically integrate the two plasmids, we opted for DNA transposons, specifically, the PiggyBac system^22^, and the Sleeping Beauty system^23^. We also decided to genomically integrate a constitutively expressed nuclear mCherry protein to facilitate the identification of cell nuclei for imaging and fluorescent experiments. For this integration, we generated a third plasmid, as shown in **Figure 3E**, in which we cloned an H2B-tagged mCherry protein under the control of the weak PGK promoter and inserted it into a vector containing Tol2 repeats for the mT2TP transposase^24^. Our integration strategy involved positive selection of integrated cells through Fluorescence Activated Cell Sorting and antibiotic resistance, as illustrated in **Figure 3B**. We first stably integrated the nuclear mCherry protein by transient transfection in Hek293T cells of the two plasmids in **Figure 2E**, one encoding for the cargo to be integrated (the nuclear mCherry) and the other for the mT2TP transposase. We then sorted for the red-positive cells three times to obtain a uniform cell population (**Figure 3B and Supplementary Fig. 3A**). The next step involved transient transfecting of the red-positive cells with the two plasmids encoding for the CRISPRaTOR together with additional two plasmids encoding for the Sleeping Beauty and the PiggyBac transposases as shown in **Figure 3B, C, D**. Antibiotic selection by puromycin followed by sorting of green fluorescent cells was then used to select for cells stably integrating the CRISPRaTOR (**Fig. 3B** and **Supplementary Fig. 3B**). Indeed, as DD-rtTA-EGFP and Puromycin resistance are expressed in the same plasmid under the control of the gRNA-inducible 7B_pMin promoter, while the transactivator VPR-dCas9-N8 together with the cognate guide RNA (gRNA_B) is expressed on the other plasmid, only cells that have been transfected with both plasmids will be positively selected by puromycin and will pass cell sorting for the green fluorescence.

The resulting cell line can be used to test both the closed-loop CRISPRaTOR against the *Open Loop* (OL) configuration. Indeed, growing the cells in the absence of Doxycycline “opens the loop” as the antiCRISPR is not expressed, whereas growing cells in the presence of Doxycycline “closes the loop”. After constructing the stable cell line, we first assessed the functionality of the CRISPRaTOR in response to saturating concentrations of Shield1 and Doxycycline. As illustrated in **Supp. Figure 3C**, in the absence of both Doxycycline and Shield1, the cells displayed no green fluorescence. Upon addition of Shield1, the population exhibited maximal green fluorescent intensity. Conversely, in the presence of both Shield1 and Doxycycline, cells reached an intermediate level of green fluorescence intensity. This is to be expected, since in this condition the rtTA protein binds to the pTRE3G promoter and drives expression of the antiCRISPR AcrIIA4. This protein then sequesters part of the VPR-dCas9 thus reducing transcription from the 7B_pMin promoter of the DD-rtTA-EGFP construct leading to a decrease in green fluorescence.

Next, we aimed at assessing the ability of the CRISPRaTOR in achieving RPA in the stable cell line, that is in maintaining the level of DD-rtTA-EGFP protein stable against changes in Shield1 concentration. To this end, we employed a High Content Screening platform comprising an Opera Phenix Imaging to quantify green fluorescence across different combinations of Doxycycline and Shield1 concentrations (**Figure 4A**). As summarised in **Figure 4B**, we tested the CRISPRaTOR at six fixed concentrations of Doxycycline from 100 ng/ml to 1000 ng/ml (Closed Loop, CL). For each fixed concentration of doxycycline, we tested 12 different concentrations of Shield 1 ranging from 0 nM to 1000 nM. As a negative control, we tested the cells in the absence of Doxycycline, thus preventing DD-rtTA-EGFP from binding to the pTRE3G promoter and expressing AcrIIA4 to close the feedback loop (Open Loop, OL). Results in **Figure 4B** show that in the Open Loop configuration, decreasing Shield1 concentration results in a proportional decrease in green fluorescence. We confirmed that this effect was specific for the green fluorescence, as the mCherry fluorescence remained constant, as shown in **Supplementary Figure 4A**. On the contrary, in the presence of Doxycycline, the CRISPRaTOR can maintain green fluorescence intensity unchanged over a larger range of Shield1 concentrations. In **Figure 4D**, the relative green fluorescence values in **Figure 4B** are reported as a bar plot for only two conditions: a fixed Doxycycline concentration of 100 ng/ml (blue bars), or without Doxycycline (green bars). It can be observed that in the presence of Doxycycline, even when Shield1 concentration is reduced by 10-fold from 1000 nM to 100 nM, green fluorescence is unchanged. This is, however, not the case in the absence of doxycycline (Open Loop), where fluorescence is reduced by about 20%. Moreover, even a 100-fold reduction in Shield 1 (10 nM) results in less than 20% reduction in fluorescence in the presence of doxycycline (Closed Loop), but in more than a 40% reduction in the absence of doxycycline (Open Loop). As a control, we also measured the mCherry fluorescence in **Supplementary Figure 4B**. We also report raw data of EGFP and mCherry fluorescence (**Supplementary Fig. 4C, D**).

In this work, we present the CRISPRaTOR, a protein-protein-based biomolecular circuit for precise control of gene expression in mammalian cells. We demonstrate that the CRISPRaTOR exhibits Robust Perfect Adaptation, as it can maintain expression of a protein of interest in the face of changes in protein degradation and in plasmid copy number. Importantly, we also show that RPA holds not only in transient transfection, but also in cells genomically integrated with the CRISPRaTOR, thus paving the way for practical applications in biotechnology that require stable cell lines.

Recently, two other implementations of AIC controllers in mammalian cells have been reported, one based on sense-antisense RNA^13^ and the other on protein split-inteins^14^. In the first implementation, an antisense RNA is produced under the control of a promoter activated by a specific transcription factor (tTA), which then binds to the tTA mRNA, creating a negative feedback loop. Despite being very versatile, this implementation has some drawbacks as it can trigger the Integrated Stress Response in the cell because of the formation of double-stranded RNA. Moreover, it can be applied only when the mRNA to be controlled is stable^13^. The second implementation uses engineered proteins with split inteins for the key sequestration reaction by exploiting a protein splicing reaction. The splicing deactivates the inteins but preserves the functions of the proteins involved, hence also allowing the implementation of more sophisticated Proportional-Integral (PI) controllers. The split-inteins implementation of the AIC however is more difficult to adapt to the control of endogenous proteins, as the engineering of proteins with inteins requires careful consideration of intein selection, insertion sites, and its effect on protein folding. In both implementations of the AIC, the authors demonstrated Robust Perfect Adaptation (RPA) only by transient transfection.

The CRISPRaTOR is a complementary implementation of the AIC in mammalian cells. It exploits the CRISPR-antiCRISPR sequestration reaction that can be easily generalised to control any endogenous transcription factor by: (1) designing the proper guide-RNA to direct VPR-dCas9 to the endogenous promoter, and (2) engineering a synthetic promoter responsive to the endogenous transcription factor to drive the expression of the anti-CRISPR. Both steps can be routinely performed with a very high success. One of the limitations of the current implementation is the large genomic size (about 15Kb) that may hinder the delivery in primary cells or tissues where viral vectors are necessary.

Finally, we generated a cell line stably integrated with the CRISPRaTOR that exhibits RPA. This cell line can be used to express any protein of interest under the control of the pTRE3G promoter guaranteeing its robust expression over time.

## AUTHOR INFORMATION

Corresponding Authors

* ORCID

Diego di Bernardo:

## Author contributions

A.M. and D.d.B. designed the research; A.M. designed, built, and experimentally validated circuits, carried out experiments and performed data analysis; V.F. and F.R. helped to perform experiments. A.M. and D.d.B wrote the paper. All authors contributed to review and editing; D.d.B. supervised the project and secured funding.

## Notes

The authors declare no competing financial interest.

## ACKNOWLEDGMENTS

We thank Luigi Ferrante from the Fluorescence Activated Cell Sorting facility for his help in the construction of the stable cell line. We also thank Sandro Montefusco and Antonella Capuozzo from the High Content Screening facility run by Prof. Diego Medina, for their support in high content imaging and fluorescence quantification experiments. This work was supported by Fondazione Telethon, by University of Naples Federico II and Compagnia di San Paolo - Programme STAR Plus. A.M. was supported by a fellowship from the European School of Moleculer Medicine (SEMM).

## Methods

### Plasmid Construction

Most of the plasmids were constructed using the Golden-Gate based EMMA assembly kit^25^, following authors’ protocol^26^, and Gibson assembly method^27^. All the sequences regarding N7 and N8 coiled coils^17^, used to modify the CRISPR-antiCRISPR system, AcrIIA4^14^ and FKBP-derived DD^28^, gRNA inducible constructs with 7 binding sites for gRNA B^18^, gRNAs sequences^18^, VPR-dCas9^29^, PiggyBac^22^, SleepingBeauty^23^ and Tol2^24^, have all been taken from published papers. The Tet-On®3G system, comprising the TRE3G promoter and the rtTA protein, has been acquired from Takara Bio.

### Cell culture and transfection

The Hek293T cell line (ATCC) was cultured in DMEM Gluta-max (Gibco) supplemented with 10% Tet-Free Fetal Bovine Serum (Euroclone) and 1% Penicillin-Streptomycin (Euroclone). Cells have been kept at 37°C in a 5% CO2 environment. For luciferase experiments, 2x10^4^ Hek293T cells per well were seeded in CoStar White 96-well plates (Corning) to perform standard transfection, while 4,5x10^4^ were seeded when performing reverse transfection. For Flow Cytometry assay the same numbers of cells have been seeded in 96-well cell culture plates (Corning). After 18 hours of seedling, for standard transfection, or immediately after seedling, for reverse transfection, cells have been transfected using a home-made solution of PEI (MW 25000, Polysciences, stock concentration 0,324 mg/ml, pH 7.5) using 250ng of DNA per well.

### Luciferase assay

To normalize reporter values to transfection efficiency, 10ng of pRL-TK (encoding for Renilla Luciferase) have been used for each experiment. The cells were collected 48 hours after transfection and lysed with 5X Passive Lysis Buffer (Biotin) diluted in water. Firefly Luciferase and Renilla Luciferase expression were measured using the Dual Luciferase Assay (Promega) on a Glomax Explorer plate reader (Promega). Firefly Luciferase Arbitrary Units (Luciferase [A.U.]) were calculated by normalizing each sample’s Firefly Luciferase activity to the constitutive Renilla activity detected in the same sample. Relative Luciferase [A.U.] values have been obtained by dividing each sample’s Luciferase [A.U.] by the Luciferase [A.U.] of the sample treated with the maximum dose of Shield1 used, 1000 nM. For the RPA experiments, a Firefly Luciferase with two destabilization sequences has been used (Luc2CP, Promega).

### Cell Sorting

For integration of the vector containing H2B-mCherry, Hek293 were seeded in a 6 well plate and transfected with the vector to integrate and the mT2TP transposase with a 2:1 molar ratio. Following transfection and expansion, cells underwent three rounds of sorting for mCherry-positive cells, using a BD FACS Aria III Cell Sorting System (Becton Dickinson). For integration of the CRISPRaTOR, H2B-mCherry containing cells were seeded in a 6 well plate and transfected with the CRISPRaTOR-codifying vectors and their respective transposases in a 3:1 molar ratio. Following 2 weeks of Puromycin selection (1,5 μg/ml), the mCherry-EGFP double positive cell were selected through cells sorting, using a BD FACS Aria III Cell Sorting System (Becton Dickinson).

### Flow cytometry

Cells were collected 48 hours after treatment, washed and resuspended with PBS (Euroclone). Flow Cytometry analysis was carried out using an Accuri C6+ (BD Biosciences), analyzing 10000 cells for each sample. A 488-nm laser with a 670 nm LP filter was used to excite and detect mCherry fluorescence, while a 488-nm laser with a 533/30 nm filter was used to excite and detect EGFP fluorescence.

### Drug treatment

Cells were treated with Doxycycline (Clontech) and/or Shield1 (MedCehmExpress) immediately before transfection. For High Content Screening and Flow Cytometry experiments, cells were treated 18 hours after seedling. Doxycycline was dissolved in H2O, while Shield1 was dissolved in DMSO. For transient transfection experiments, drugs’ concentrations are referred to the medium volume before adding the transfection mix.

### High Content Screening and Fluorescence Quantification

For High Content Screening experiments, 1x10^4^ (Hek293T) cells per well were seeded in PhenoPlate 96-well plates (Perkin Elmer) and, the day after, treated with Doxycycline and Shield1. Forty-eight hours after treatment, cells were fixed in 4% PFA and imaged with the Opera Phenix High Content Screening System (Perkin Elmer), acquiring at least 10 images per well. Cells’ nuclei have been identified through mCherry fluorescence. Fluorescent signals (mCherry and EGFP) were quantified using a custom script developed on Signals Image Artists (Perkin Elmer).

